# Improved the Protein Complex Prediction with Protein Language Models

**DOI:** 10.1101/2022.09.15.508065

**Authors:** Bo Chen, Ziwei Xie, Jiezhong Qiu, Zhaofeng Ye, Jinbo Xu, Jie Tang

**Affiliations:** Department of Computer Science and Technology, Tsinghua University, Beijing, China; Toyota Technological Institute at Chicago, Chicago, IL 60637, USA; Tencent, Shenzhen, China

**Keywords:** Protein Complex Structure Prediction, Protein Language Model

## Abstract

AlphaFold-Multimer has greatly improved protein complex structure prediction, but its accuracy also depends on the quality of the multiple sequence alignment (MSA) formed by the interacting homologs (i.e., interologs) of the complex under prediction. Here we propose a novel method, denoted as ESMPair, that can identify interologs of a complex by making use of protein language models (PLMs). We show that ESMPair can generate better interologs than the default MSA generation method in AlphaFold-Multimer. Our method results in better complex structure prediction than AlphaFold-Multimer by a large margin (+10.7% in terms of the Top-5 best DockQ), especially when the predicted complex structures have low confidence. We further show that by combining several MSA generation methods, we may yield even better complex structure prediction accuracy than Alphafold-Multimer (+22% in terms of the Top-5 best DockQ). We systematically analyze the impact factors of our algorithm and find out the diversity of MSA of interologs significantly affects the prediction accuracy. Moreover, we show that ESMPair performs particularly well on complexes in eucaryotes.

## 1 Introduction

Most proteins function in a form of protein complexes[1–5]. Consequently, obtaining accurate protein complex structures is vital to understanding how a protein functions at the atom level. Experimental methods, such as X-ray crystallography and cryo-electron microscopy, are costly and low-throughput, and require intensive efforts to prepare samples for structure determination. The computational methods, termed as protein complex prediction (PCP) or protein-protein docking, is an attractive alternative for solving complex structures. PCP takes sequences and/or the unbound structures of individual protein chains as inputs and then predicts the bound complex structures. Traditional computational methods often rely on the global search paradigm, such as fast-Fourier transform based methods like ClusPro [6], PIPER [7], and ZDOCK [8] and Monte Carlo sampling-based methods like RosettaDock [9], have been widely used in practice. These methods exhaustively search the conformation space of a complex, and optimize score functions to obtain the final structures. Since the conformation space is large, these methods have to make restrictive constraints on the search space in order to obtain results within a reasonable amount of time. Typical constraints include reducing the search resolutions, making the input monomers rigid bodies, and using score functions that can be quickly evaluated [7, 8]. As a result, global search methods have relatively low prediction accuracy and are used with more computationally intensive local refinement methods to obtain higher resolution predictions [10]. To date, PCP is still a fundamental and longstanding challenge in computational structural biology [11, 12]. Various methods have been proposed for PCP, but with limited accuracy. When only sequences are given as inputs, PCP is even harder because the unbound structures of individual chains and auxiliary information on the complex interfaces are unavailable.

In the last decades, deep learning has enabled substantial progress in quite a few computational structural biology tasks, such as protein contact [13–15], tertiary structure prediction [16–18], and cryo-electron microscopy structure determination [19, 20]. Among these, co-evolution analysis based contact prediction [18, 21, 22] and structure prediction [23, 24] have made substantial progress and demonstrated state-of-the-act accuracy for monomers. These methods utilize the co-evolutionary information hidden in MSA to infer inter-residue interactions or three-dimensional structures of the targets. AlphaFold2 is the representative method, which has showed unparalleled accuracy in CASP14 [16]. Recently, AlphaFold-Multimer [25], a derived version of AlphaFold2 for multimers, significantly outperforms prior protein complex prediction systems [6, 23, 24]. However, compared to the accuracy of AlphaFold2 [16] on folding monomers, the accuracy of AlphaFold-Multimer on predicting the protein complex structures is far from satisfactory. Its success rate is around 70% and the mean DockQ score is around 0.6 (medium quality judged by DockQ) [24]. The most important input feature to AlphaFoldMultimer is the multiple sequence alignment (MSA) [23, 24]. Compared with AlphaFold2 [16] that takes the MSA of a single protein as the input, AlphaFoldMultimer needs to build an MSA of interologs for protein complex structure prediction. However, how to construct such an MSA is still an open problem for heteromers. It requires the identification of interacting homologs in the MSAs of constituent single chains, which may be challenging since one species may have multiple sequences similar to the target sequence (paralogs). Several algorithms have been proposed to identify putative interologs from genome data, such as profiling co-evolved genes [26], and comparing phylogenetic trees[27]. Genome co-localization and species information are two commonly used heuristics to form interologs for co-evolution-based complex contact and structure prediction [25, 28]. Genome co-localization is based on the observation that, in bacteria, many interacting genes are coded in operons [29, 30] and are cotranscribed to perform their functions. However, this rule does not perform well for complexes in eukaryotes with a large number of paralogs, since it becomes more difficult to disambiguate correct interologs [28, 31]. The other phylogeny-based method for identifying interologs is first proposed in ComplexContact [28] and later similar ideas are adopted by AlphaFold-Multimer. This method first identifies groups of paralogs (sequences of the same species) from the MSA of each chain, then ranks the paralogs based on their sequence similarity to their corresponding primary chain, and last pairs sequences of the same species and with the same rank together. However, they are all hand-crafted approaches which merely take effects on the specific domains. In this paper, we instead investigate general and automatic algorithms for constructing MSAs of interologs for heterodimers effectively.

Representation learning via pre-training techniques has been prevailing in different applications [34–37]. Inspired by this, protein language models [38–40] (PLMs) have surged as the main regime for protein representation learning built on a large amount of protein sequences, which benefits downstream tasks, such as contact prediction [15, 39], remote homology detection [41, 42] and mutation effect prediction[43]. PLMs can comprehensively capture the biological constraints and co-evolutionary information encoded in the sequence, which is a plausible interpretation for their impressive performance on various downstream tasks than canonical methods relying on dedicated hand-crafted traits. To this, a natural question arises: ***Can we leverage the co-evolutionary information featured by PLMs to build effective interologs?***

In this paper, we mainly focus on *ab-initio* protein complex structure prediction, i.e., predicting the complex structure without prior information on the binding interfaces of the target complex. To our best knowledge, we are the first to propose a simple yet effective MSA pairing algorithm that uses the immediate output from protein language models to form joint MSAs, i.e., MSA of interologs. In particular, we leverage column-wise attention scores from ESM-MSA-1b [39] to identify and pair homologs from MSAs of constituent single chains, coined as ESMPair. We conduct extensive experiments on three test sets, i.e., pConf70, pConf80, and DockQ49. Compared with previous methods, ESMPair achieves state-of-the-art structure prediction accuracy on heterodimers (+10.7%, +7.3%, and +3.7% in terms of the Top-5 best DockQ score over AlphaFold-Multimer on three test sets, respectively). Moreover, we find out that the ensemble strategies, which combine ESMPair with other MSA pairing methods, significantly improve the structure prediction accuracy over the standard single strategy. We further analyze the performance of complexes from eukaryotes, bacteria, and archaea, and find out ESMPair performs the best on eukaryotes for which identifying interologs is quite difficult [28, 31]. Most strikingly, on a few targets where one of the constituent chains is from eukaryotes while the other is from bacteria, ESMPair considerably outperforms other baselines (+25% in overall performance over AlphaFold-Multimer), which strongly demonstrates that the PLM-enhanced MSA pairing method is robust for targets from different superkingdoms. Then we exposit that the diversity of interologs has a significant positive correlation with the prediction accuracy. Lastly, we explore other approaches that utilize the output of ESM-MSA-1b. For example, we take the cosine-similarity score between the sequence embeddings as the metric to build interologs, which performs on par with the default protocol used in Alphafold-Multimer. Generally, ESMPair is the first simple yet effective algorithm that incorporates the strength of PLMs into tackling the issues of identifying MSA of interologs. We believe ESMPair will facilitate the fields of protein structure prediction which highly resorts to the co-evolution information hidden in MSA.

## 2 Results

In this section, we first briefly outline the framework of ESMPair for protein complex prediction (Section 2.1). Then, we discuss our proposed methods obtain better complex prediction accuracy than previous MSA pairing methods (Section 2.2). We find out the ensemble strategy showcase the excellent performance that the default single strategy (Section 2.3). We further quantitatively analyze several key factors and hyperparameters that may impact the performance of our method, and also explore the capability of different measurements to distinguish acceptable predictions from unacceptable ones (Section 2.4).

### 2.1 ESMPair overview

The overall framework of ESMPair is illustrated in Fig. 1 with the details in Methods. In complex structure prediction, predictors such as AlphaFoldMultimer make use of inter-chain co-evolutionary signals by pairing sequences between MSA of constituent single chains of the query complex. Formally, given a query heterodimer, we obtain individual MSAs of its two constituent chains, denoted as 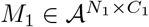 and 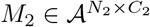, where *A* is the alphabet used by PLM, *N*_1_ and *N*_2_ are the number of the sequences in MSAs *M*_1_ and *M*_2_, and *C*_1_ and *C*_2_ are the sequence length. The MSA pairing pipeline aims at designing a matching or an injection *π* : [*N*_1_] → [*N*_2_] between MSAs from each chain to build the MSA of interologs, dubbed as 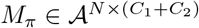, where *N* is the number of the sequence in the joint MSA. In practice, the MSA of interologs *M*_*π*_ is a collection of the concatenated sequence *{*concat(*M*_1_[*i*], *M*_2_[*π*(*i*)]) : *i ∈ 𝒫}*, where *𝒫* is the indices of the sequences from *M*_1_ that can be paired with any sequences from *M*_2_ according to the matching pattern *π*. Then MSA of interologs is taken by predictors as input to predict the structure of the query heterodimer. Our aim is to leverage the superiority of PLMs to explore an effective matching strategy *π* that facilities the protein complex structure prediction.

**Fig. 1.**
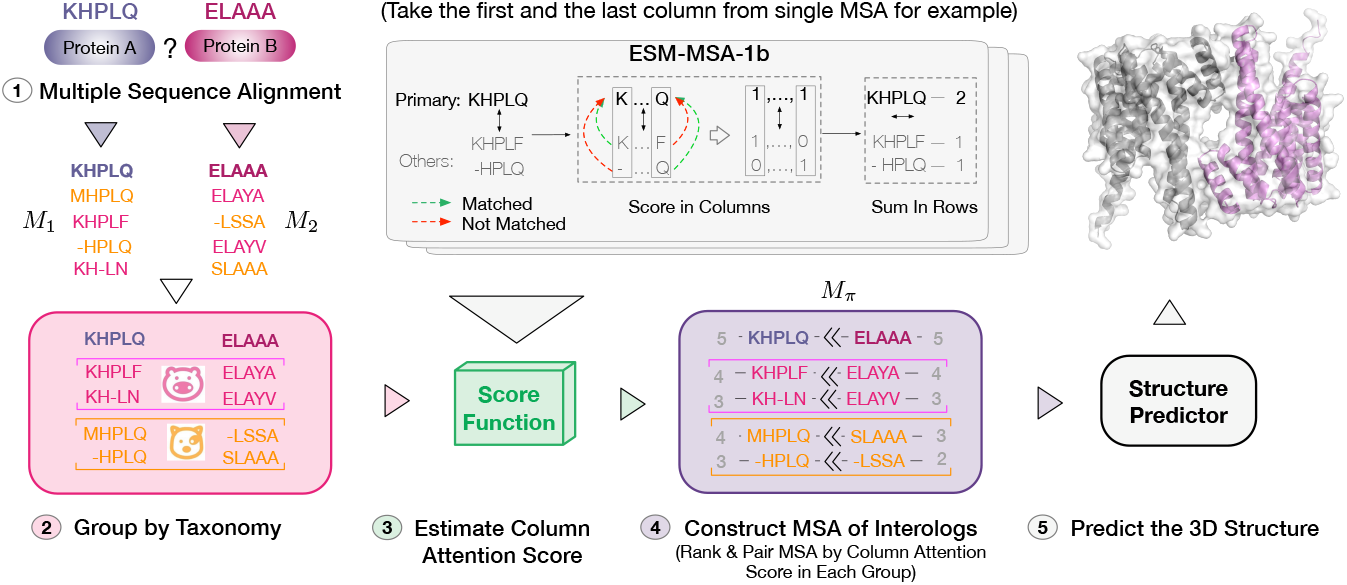
Schematic illustration of ESMPair that builds interologs as the input to AlphaFold-Multimer. Given a pair of query sequences as input: 1) we first search the UniProt database [32] with JackHMMER [33] to generate the MSA for each query sequence, 2) sequences of the same taxonomy rank are grouped into the same cluster, 3) ESM-MSA-1b is applied to estimate the column attention score (ColAttn score) between each sequence homolog of MSA with the query sequence. 4) One interolog is obtained by directly concatenating two matched sequence homologs. We match two sequence homologs of the same taxonomy group with similar attention scores from the two query sequences, 5) AlphaFold-Multimer takes the interolog MSA as input to predict the complex structure.

### 2.2 ESMPair outperforms other MSA pairing methods on heterodimer predictions

#### Overall evaluation

For each test target we predict five 3D structures using Alphafold-Multimer’s 5 models and then report the average of Top-*k* (k=1, 5) Best DockQ score of the predicted structures and the corresponding success rate (SR) in Table 1. Our method outperforms the other methods. To be specific, our method outperforms the AF-Multimer’s default MSA pairing strategy on all three test sets (0.259 vs. 0.234 on pConf70, 0.423 vs. 0.406 on pConf80, and 0.265 vs. 0.242 on Quality49, in term of Top-5 DockQ score). Our experimental results confirm that our proposed column-wise attention based MSA pairing method, denoted as ESMPair, is better than 1) the sequence similarity-based method used in AF-Multimer, and 2) the cosine similarity-based method based on the mixed noisy residue embedding, i.e., ESMPair(InterLocalCos), ESMPair(InterGlobalCos), and ESMPair(IntraCos). Hereinafter, we abbreviate them as IntraLocalCos, InterGlobalCos, and InterCos.

**Table 1.**
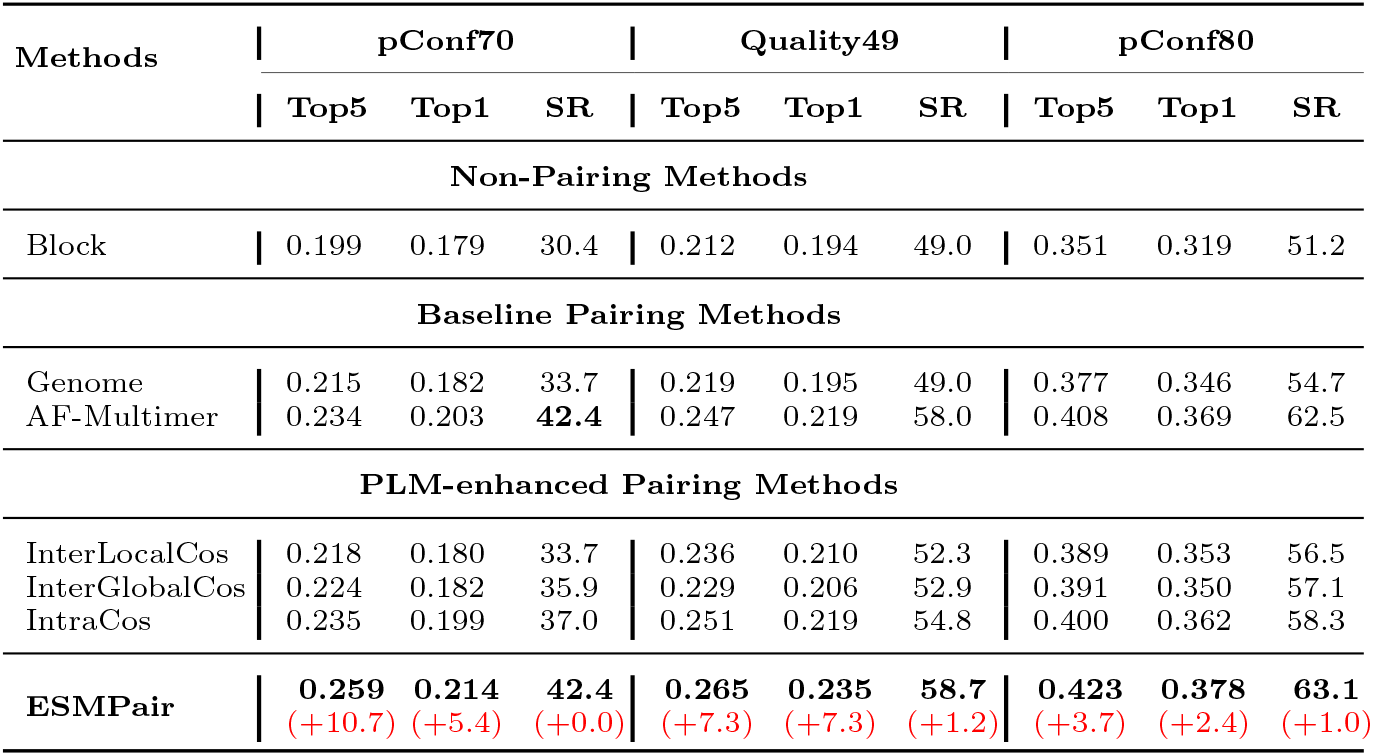
DockQ scores and success rate of PLM-enhanced pairing methods and baselines. We report the average of Top-5 Best DockQ score, Top-1 Best DockQ score, and Success Rate (DockQ ≥0.23) (%) on pConf70, Quality49, and pConf80 test sets. For one test target, we predicted 5 different structures using the five AlphaFold-Multimer models. Subscript in red represents the performance gain of our method over the default MSA pairing strategy in Alphafold-Multimer (%).

| Methods | pConf70 |  |  | Quality49 |  |  | pConf80 |  |  |
| --- | --- | --- | --- | --- | --- | --- | --- | --- | --- |
|  | Top5 | Top1 | SR | Top5 | Top1 | SR | Top5 | Top1 | SR |
| **Non-Pairing Methods** |  |  |  |  |  |  |  |  |  |
| Block | 0.199 | 0.179 | 30.4 | 0.212 | 0.194 | 49.0 | 0.351 | 0.319 | 51.2 |
| **Baseline Pairing Methods** |  |  |  |  |  |  |  |  |  |
| Genome | 0.215 | 0.182 | 33.7 | 0.219 | 0.195 | 49.0 | 0.377 | 0.346 | 54.7 |
| AF-Multimer | 0.234 | 0.203 | **42.4** | 0.247 | 0.219 | 58.0 | 0.408 | 0.369 | 62.5 |
| **PLM-enhanced Pairing Methods** |  |  |  |  |  |  |  |  |  |
| InterLocalCos | 0.218 | 0.180 | 33.7 | 0.236 | 0.210 | 52.3 | 0.389 | 0.353 | 56.5 |
| InterGlobalCos | 0.224 | 0.182 | 35.9 | 0.229 | 0.206 | 52.9 | 0.391 | 0.350 | 57.1 |
| IntraCos | 0.235 | 0.199 | 37.0 | 0.251 | 0.219 | 54.8 | 0.400 | 0.362 | 58.3 |
| **ESMPair** | **0.259**<br>(+10.7) | **0.214**<br>(+5.4) | **42.4**<br>(+0.0) | **0.265**<br>(+7.3) | **0.235**<br>(+7.3) | **58.7**<br>(+1.2) | **0.423**<br>(+3.7) | **0.378**<br>(+2.4) | **63.1**<br>(+1.0) |

Among all the MSA pairing methods, block diagonalization performs the worst (−30% compared with ESMPair in terms of the average of Top-5 best DockQ). The result indicates that the inter-chain co-evolutionary information helps with complex structure prediction. Among MSA pairing baselines, AF-Muiltmer surpasses genetic co-localization by a large margin (+12.8% Top-5 DockQ). All the proposed PLM-enhanced pairing methods substantially outperform the block diagonalization and the genetic-based methods. Even though AF-Multimer may have overly optimistic performance using the default pairing method since the training MSAs are built using it, IntraCos MSA pairing method performs on a par with AF-Multimer, and ESMPair further exceeds it by a large margin (+4.2~10.7% Top-5 DockQ score over three test sets).

#### Intra-ranking methods are superior to inter-ranking ones both in effectiveness and scalability

From Table. 1, we can also see inter-ranking methods like InterLocalCos and InterGlobalCos underperform the intra-ranking ones, i.e. IntraCos and ESMPair. We speculate that as ESM-MSA-1b pre-trains in the monomer data, it fails to directly capture the underlying correlations across the constituent chains in the complex. Besides heterodimers, when it extends to predict the structure of multimer with more than two chains, intra-ranking strategies are the self-contained methods that only need to rank the MSAs in each single chain, and then match MSA of the same rank with other chains to build effective interologs with time complexity of *O*(*N*), where *N* is the depth of MSA. While the inter-pairing strategies suffer from the exponential growth of combinations with increasing interacting chains with the time complexity *O*(*N* ^*r*^), where *r* is the number of chains in the multimer. Thus, intra-ranking methods are more time-efficient and scalable than inter-ranking ones.

#### ESMPair performs better on low pConf targets

As shown in Table. 1, the performance gap between ESMPair and AF-Multimer becomes narrower on pConf80 than on pConf70, with improvement ratio from 3.7% to 10.7%. To take an in-depth analysis, we quantitatively analyze the correlations between the predicted confidence score (pConf) estimated by AF-Multimer and the performance gap of the average of Top-5 Best DockQ score between ESM-Pair and AF-Multimer on Quality49, as illustrated in Fig. 3(e-f). The relative improvement is negatively correlated (Pearson Correlation Coefficient is −0.49) with the predicted confidence score. When pConf is less than 0.2, the relative improvements even achieve 100%, while when pConf is more than 0.8, ESMPair performs nearly on par with AF-Muiltimer. This is because AF-Multimer can do well on a relatively easier target, it is very challenging to further improve it.

#### ESMPair has the higher prediction accuracy on eucaryote targets

We further compare the DockQ distribution of ESMPair, AF-Multimer, and Genome on three kingdoms, i.e. Eucaryote, Bacteria, and Eucaryote&Bacteria, as shown in Fig. 3(a-d), we can see that ESMPair rivals the other two MSA pairing methods on the Eucaryotes data by a large margin (0.420 for ESMPair, 0.402 for AF-Multimer, and 0.369 for Genome on the overall data). As we all know that it is notoriously different to identify homologous protein sequences for the Eucaryotes data, ESMPair has a desirable property to build effective interologs on the Eucaryotes. While in the Bacteria data, three strategies have similar performance (around 0.35 on the whole data). Most strikingly, we find ESMPair has an extraordinary performance on the Euca.&Bact. data over the other two methods (0.394 for ESMPair, 0.314 for AF-Multimer, and 0.277 for Genome on the overall data). We further check the performance gap for each target from the Euca.&Bact. data. ESMPair performs significantly better on the three out of six targets, 0.443 (ESMPair) versus 0.013 (AF-Multimer) on 5D6J, 0.289 versus 0.201 on 6B03, and 0.864 versus 0.854 on 7AYE. Besides, ESMPair performs on par with AF-Multimer on the other three targets. These results shed light on the robustness of protein language models. As PLMs are pre-trained on billions of protein data [38–40], it can break the bottleneck that other hand-crafted MSA pairing methods, such as genetic-based methods, phylogeny-based methods, etc, which merely take effect in the specific domain. While our proposed PLMs-enhanced methods can identify the co-evolutionary signals effectively to build MSA of interologs across different superkingdoms.

#### ESMPair outperforms AF-Multimer on the most of newly-released targets

We further select 74 targets that AF-Multimer does not train on [25], i.e., the targets whose release date is later than 2018-4-30 from the test dataset. Then we compare the performance of predicted structures on these targets between ESMPair and AF-Multimer in Fig. 2. From Fig. 2(a), ESMPair outperforms AF-Multimer on the most of targets (57%) with a relative larger performance gap, while AF-Multimer outperforms ESMPair on fewer targets (35%) with a relative lower gap. We further plot the distributions between interface RMSD and ligand RMSD of predicted structures via ESMPair and AF-Multimer in Fig. 2(b). The holistic distributions predicted by ESMPair are closer to the origin of coordinates than that predicted by AF-Multimer, which strongly proved ESMPair is superior to AF-Multimer on the predictions of newly-released targets.

**Fig. 2.**
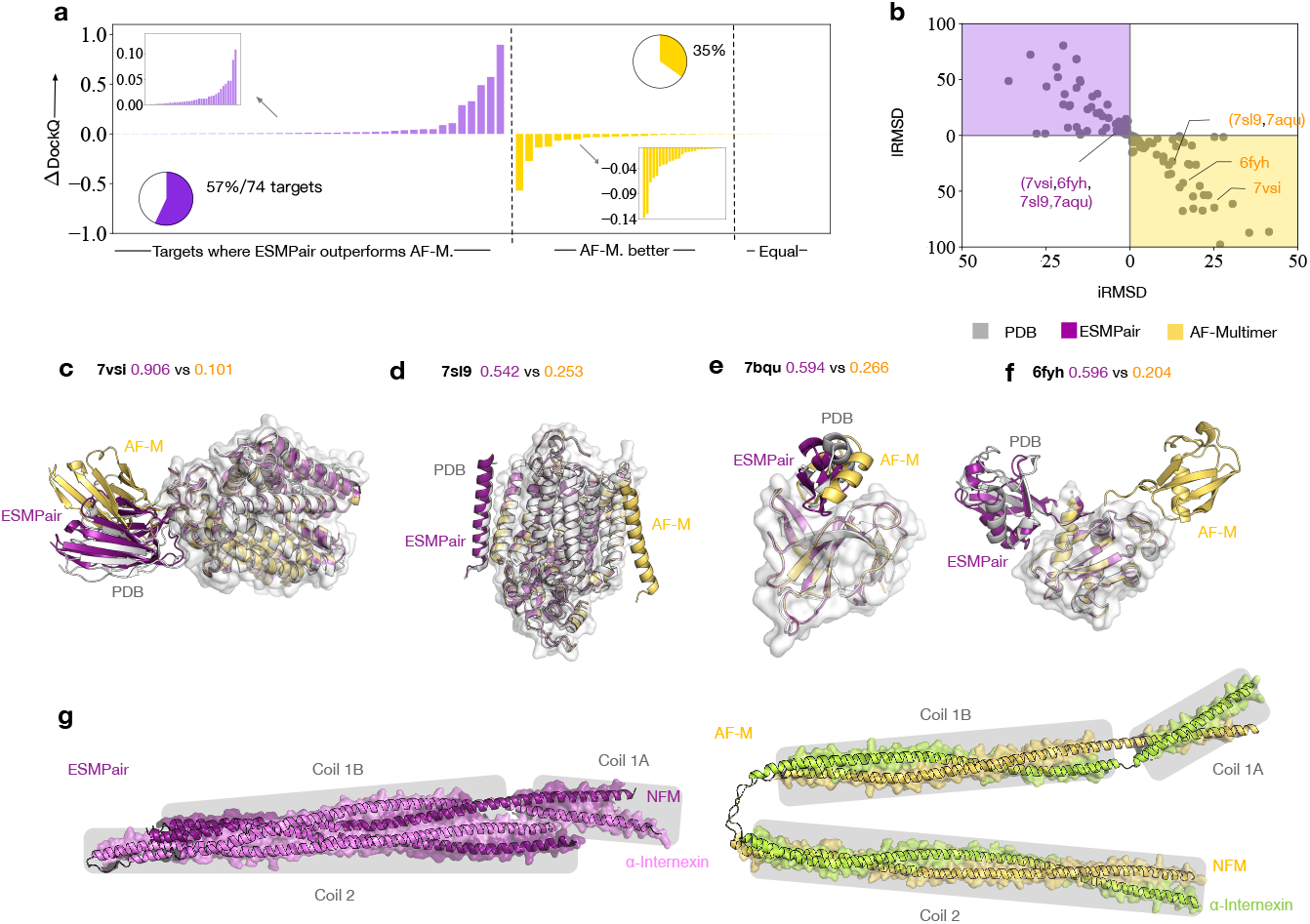
Comparing ESMPair and AF-Multimer predictions on newly-release targets (a-f) and unresolved cases (g). **a-f**, The comparisons are made on the 74 targets whose release data is later on 2018-4-30. **a**, The bar charts show the relative performance gap between ESMPair and AF-Multimer on three categories: ESMPair outperforms AF-M.; AF-M. outperforms ESMPair; Equal performance. **b**, The interface and ligand RMSD distributions of predicted stuctures via ESMPair (Purple) and AF-M (Yellow). **c-f**, Four cases are further visualized. Among this, the ligand orientation are wrongly-predicted via AF-M. on 7VSI and 7AQU, while the binding site are wrongly-predicted by AF-M. on 7SL9 and 6FYH. **g**, The intermediate filament protein NFM-INA heterodimer structure predicted via ESMPair shows a four-helix bundle. The gray boxes show the interacting motifs of coil 1A, coil 1B and coil 2 of the two proteins.

Furthermore, we show why ESMPair performs better than AF-Multimer by analysing four PDB targets, 7VSI, 7AQU, 6FYH, and 7VSI. in Fig. 2(c-f). Among these, 7VSI and 6FYH have a larger predicted iRMSD and lRMSD variance by AF-Multimer, because AF-Multimer predicts the wrong binding sites. While AF-Multimer predicts the right binding sites on 7SL9 and 7AQU that have a smaller predicted iRMSD and lRMSD variance, it unfortunately predicts the wrong ligand orientations. By contrast, our proposed ESMPair correctly predicts the binding sites on the receptor and also places the ligand in the approximately correct relative orientation.

To better illustrate the usage of ESMPair in predicting the protein complexes without known resolved 3D structures, we inspected the intermediate filament heterodimer formed between the neurofilament medium polypeptide (NFM, UniProt ID P08553) and *α*-internexin (UniProt ID P46660), which is known to form an anti-parallel four-helix bundle[44, 45]. As shown in Fig. 2(g), both ESMPair and AF-Multimer correctly predict the three binding interfaces between the coil 1A, coil 1B and coil 2 motifs from NFM and *α*-internexin. However, ESMPair predicted the two coiled coils to pack as a four-helix bundle, which is consistent with the experimental evidences, while the AF-multimer predicted the two coiled coils to be separated. This case demonstrate the potential to apply ESMPair to model unresolved protein complexes.

### 2.3 Ensemble improves the prediction accuracy

From Fig. 4 (a-d), we found that different MSA pairing methods have their own advantages, even block diagonalization performs slightly better than ESMPair on about 30% targets, which implies that they can complement each other. To verify that, we combine ten models predicted by any two of the MSA paring methods, then we report the average of Top-5 Best DockQ score, as shown in Fig. 4 (e). The ensem strategies, i.e., the yellow and purple bars, significantly outperform the corresponding single strategy, i.e., the grey bars. Specifically, the performance of intra-ensemble strategies surpass the corresponding single strategy, for example, the DockQ score of ESMPair + ESMPair is 0.269 versus 0.259 of ESMPair, which demonstrates that simply increasing the number of predictions of each model also benefits the structure prediction accuracy of each target. Among the inter-ensemble strategies, ESMPair pluses any one of the single strategy always have a better performance than the one without ESMPair, for example, the SR of ESMPair + Genome is 44.6% versus 40.4% of AF-Multimer + Genome. Finally, the ensemble of all three strategies, i.e., the purple bar, reaches the best performance with 0.285 DockQ score and 46.8% Success Rate, which motivates us that instead of merely using a single strategy to build interologs, the ensemble MSA pairing strategy may be the silver bullet to identify more effective interologs.

**Fig. 3.**
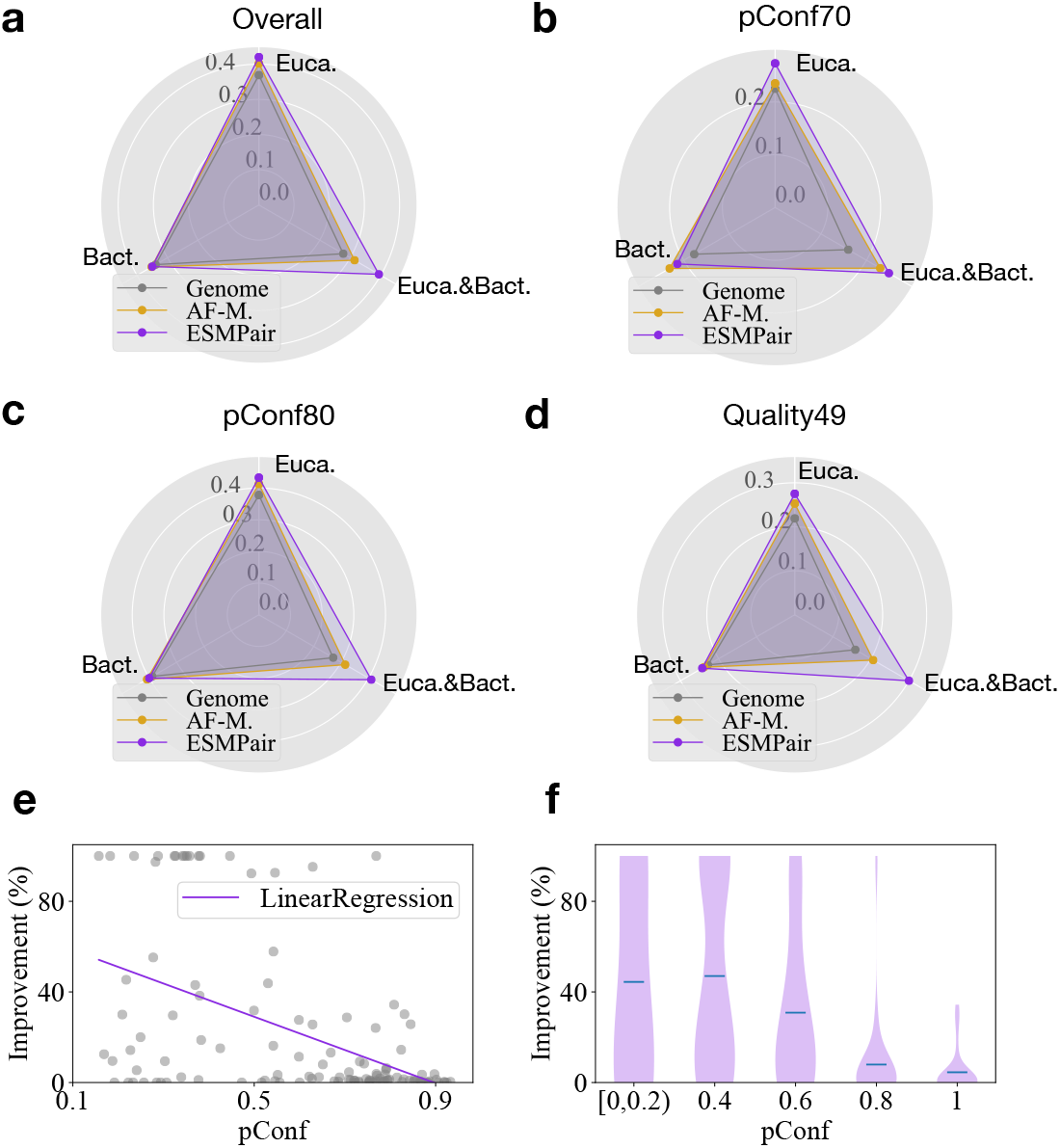
Comparison of the prediction performance on different domains. **a-d**, We compare the DockQ score among ESMPair, AF-Multimer, and Genome on Eucaryote, Bacteria, and Eucaryote&Bacteria domains. The Euca.&Bact. is a special domain means the two constituent chains in the heterodimer belong to the two domains respectively. Specifically, the heterodimers of our dataset are from Eucaryotes, Bacteria, Viruses, Archaea, Eucaryotes;Bacteria respectively. We group the data from Bateria, Viruses, and Archaea as the Bateria domain. In all test sets, ESMPair significantly outperforms other two baselines on the Eucaryote targets. **e-f**, The negative correlations between the relative improvement between ESMPair and AF-Multimer.

**Fig. 4.**
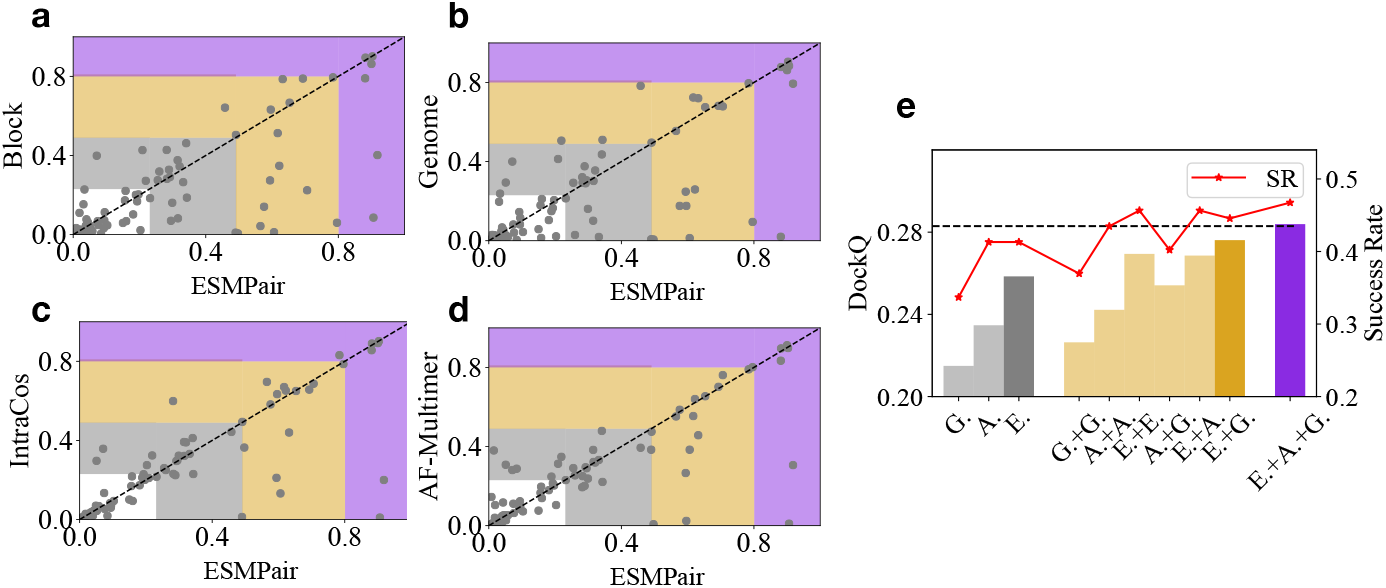
The comparisons between ESMPair and four alternative MSA pairing approaches (a-d), and various ensemble strategy (e) on the targets from pConf70. **(a-d)**, The coordinates of each point demonstrate the reported DockQ score of the target between ESMPair (x-axis) and other methods (y-axis). A point under the diagonal dash line implies ESMPair performs better than the compared method on this target. The highlight regions represent the incorrect (white), acceptable (grey), medium (yellow), and high-quality (purple) predicted models according to DockQ score. **(e)**, The grey bars represent the performance of single strategies, where G. stands for Genome, A. is for AF-Multimer, and E. is for ESMPair. ESMPair is the best with 0.259 DockQ score and 42.4% Success Rate. The yellow bars show the ensemble performance of the two strategies. Among these, ESMPair + Genome performs the best with 0.277 DockQ score with 44.6% Success Rate. The purple bar implies the best performance about the ensemble of all the three strategies with 0.285 DockQ score with 46.8% Success Rate.

### 2.4 More analytic studies of ESMPair: key factors, hyperparameters, and measurements to identify high-quality predictions

In this part, we analytically and empirically investigate the inherent properties of ESMPair. Generally, we find out the diversity of the formed MSA of interologs has a strong correlation with the performance of ESMPair. Moreover, we study the effect of different layers of ESM-MSA-1b [39] on identifying homologs. Lastly, we demonstrate the predicted confidence score output by AlphaFold-Multimer is a rational measurement to discriminate correct predictions from incorrect ones.

#### The diversity about MSA of interologs affects the predicted structure accuracy by ESMPair

We investigate the connections between the performance of ESMPair and some key factors of the formed MSA of interologs, such as the column-wise attention score (i.e., ColAttn score), the number of effective sequences within MSA measured by Meff (i.e., #Meff), the number of species (i.e., #Species), and the depth of MSA (i.e., MSA Depth). To be specific, we predict 1,689 heterodimers sampled from PDB without filtering and divide them into different regions according to the value of each factor. Notably, for ColAttn score, we average the score of each single chain in interolog, then re-scaling it in the logarithm form, and then averaging ColAttn score of all interologs from the paired MSA as the final score of the target. For #Meff, #Species, and MSA Depth, we directly calculate the corresponding statistics based on the interologs.

The correlations between DockQ score and each of above factors are illustrated in Fig. 5. #Meff, #Species, and MSA Depth have a similar trend that the predicted structure accuracy improves with the increasing of these factors. It implies that MSA with more diversity represents the more co-evolutional information that benefits structure predictions of AF-Multimer, which also meets with previous insights[39]. Moreover, the increasing ColAttn score results in the decreasing structure prediction accuracy. Considering the self-attention mechanism in the protein language model, given a sequence as the query, the self-attention mechanism aims at identifying the sequence with high homology affinity, i.e., the sequence with a high similarity score [15]. Therefore, a large ColAttn score indicates the MSA with a low #Meff, which potentially results in an inaccurate structure prediction. To justify our speculation, we explicitly characterize the dependency between ColAttn score and #Meff, as shown in Fig. 5 (e). ColAttn score has shown a negative correlation to the #Meff, with the Pearson correlation coefficient of −0.70, which elucidates that a higher ColAttn score reflects MSA with lower sequence diversity.

**Fig. 5.**
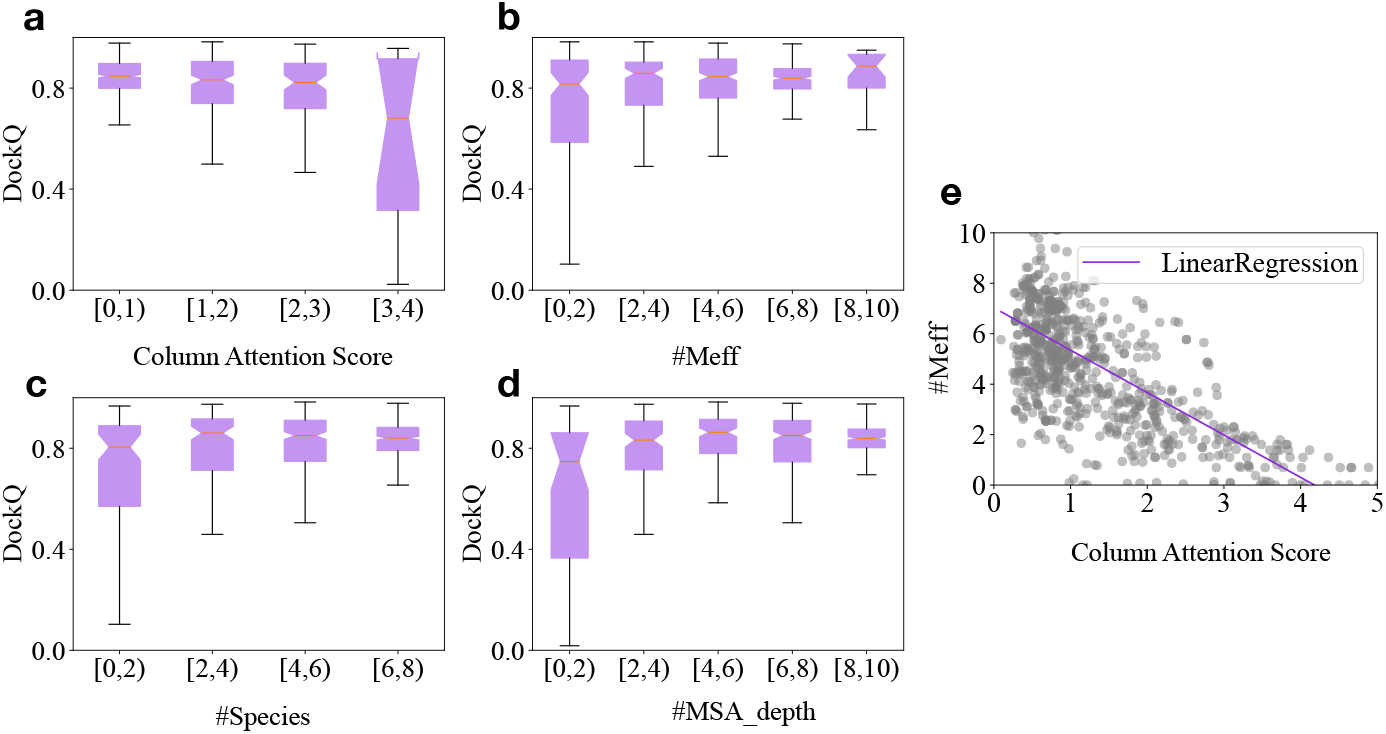
Different factors affect the performance of structure prediction. The correlations between the average of Top-5 Best DockQ score (Y-axis) and **(a)** the column attention score predicted by ESM-MSA-1b, **(b)** the number of effective sequences measured by Meff, **(c)** the number of species, and **(d)** the depth of matched MSA. **(e)** The distribution of column attention score (X-axis) and the number of effective interologs in the paired MSA (Y-axis). The red curve is the visualization of the fitted linear regression model. The Pearson correlation coefficient is about −0.70, which strongly indicates that an increasing column attention score results in the decreasing number of effective interologs.

#### ESMPair built on the last few transformer Layers has the better performance

As ESMPair leverages the column-wise attention output by ESM-MSA-1b[39] to rank and match interologs, how do the column-wise attention weight matrices by different transformer layers affect the efficacy of ESMPair? To answer this, we use the DockQ score of predicted structures as the metric to measure the quality of the input interologs built by ESMPair, as shown in Supplement Fig. A2. ESMPair that based on the attention output of layer 6 (0.258 DockQ score and 40.2% Success Rate), layer 7 (0.249 and 43.0%), and AVG (0.262 and 42.2%) perform better than other layers. Overall, the AVG aggregation of all the layers is relatively superior to others, thus we use AVG as the default setting of ESMPair. What’s more, ESMPair which built on the last few layers (6-12th) identifies homologous sequences more precisely than the former layers (1-5th). The phenomenon is consistent with the empirical insights about how to effectively fine-tune the pre-trained language models in the downstream tasks: the last few layers are the most task-specific, while the former layers encode the general knowledge of the training data[46–48], thus only aggregating latter layers may exploiting more homologous information form MSAs. We leave this in future work.

#### Predicted confidence score as an indicator to distinguish acceptable models

Practically, besides the substantial improved DockQ performance through ESMPair, it is also vital to figure out how to identify the correct models (DockQ ≥ 0.23) from incorrect ones [24]. To achieve this, we also predict all the 1,689 heterodimers via ESMPair, then we apply: 1) the predicted Confidence Score (pConf), 2) Interface pTM (ipTM), 3). predicted TM-score (pTM), and 4) the number of contacts between residues from two chains (the distance of *C*_*β*_ atoms in the residues from different chains within 8 Å) (Contacts) as the metric to rank models, as shown in Supple. Fig. A1. From Fig. 1(a), we find both pConf and ipTM are capable of distinguishing acceptable models from unacceptable ones with AUC of 0.97. pTM has a worse performance with AUC of 0.85, as pTM is used as the pessimistic predictor to measure the predicted structure accuracy of each single chain, it ignores the interactions between chains. Contacts merely count the number of interacting residues from different chains, which hardly indicates the accuracy of the predicted structure. pConf and iPTM both consider the structure in both the single chain and interfaces, which are considerate indicators to validate the quality of the predicted structure. We further quantify the interplays between pConf and DockQ score of the predicted structure, as shown in Fig. 1(b), which further confirms the strong correlations between pConf and the structure prediction accuracy.

## 3 Methods

In this part, we introduce the framework of our proposed PLMs-enhanced MSA pairing method, i.e., ESMPair. Besides, we explore other promising alternative pairing methods built on PLMs, such as InterGlobalCos, InterGlobalCos, and IntraCos. The overall framework of ESMPair is illustrated in Fig. 1.

### 3.1 The PLM-enhanced MSA pairing pipeline

Previous efforts [38–40] have confirmed that protein language models (PLMs) can characterize the co-evolutionary signals and biological structure constraints encoded in the protein sequence. Moreover, the MSA-based PLMs [15, 39] further explicitly capture the co-evolutionary information hidden in MSAs via axial attention mechanisms [49, 50]. In light of this, we adopt the state-of-the-art MSA-based PLM, i.e., ESM-MSA-1b [39], as the basis to explore how to utilize them to build rational MSA of interologs to improve the protein complex prediction based on Alphafold-Mutimer [25].

### Column Attention (ESMPair)

The column attention weight matrix, which is calculated via each column of MSA via ESM-MSA-1b, can be treated as the metric to measure pairwise similarities between aligned residues in each column. Formally, for each chain, we have the MSA *M ∈ 𝒜* ^*N×C*^. The collections of column attention matrices are denoted as {*A*_*lhc*_ *∈ ℝ*^*N×N*^ : *l ∈* [*L*], *h ∈* [*H*], *c ∈* [*C*]}, where *L* is the number of layers in PLM, *H* is the number of attention heads of each layer, and *C* is the sequence length, i.e., the number of residues of each sequence. We first symmetrize each column attention matrix, and then aggregate the symmetrized matrices along the dimension of *L, H* and *C* to obtain the pairwise similarity matrix among the sequences of MSA, denoted as *S ∈* ℝ^*N×N*^ (Eq.(1)). *S* is symmetric and its first row *S*_1_ ∈ ℝ^1*×N*^ can be viewed as measuring similarity scores between the query sequence and other sequences in the MSA,

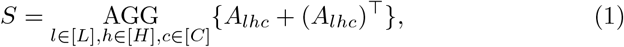

where Τ represents the transpose operation and AGG is an entry-wise aggregation operator such as entry-wise mean operation MEAN(·), sum operator SUM(·), etc. Unless otherwise specified, AGG is specified as SUM(·) in this paper.

The MSA pairing strategy is specified as follows, for a query heterodimer, we first obtain *S*_1_ of individual MSAs of constituent single chains. Then we group sequences from the MSA by their species, and rank sequences according to their similarity score of *S*_1_ in each MSA, respectively. Finally, the sequences of each MSA with the same rank in the same species group are concatenated as interologs.

#### Cosine Similarity

The cosine similarity measurement has been thoroughly explored by pre-train language models [51, 52]. Intuitively, as PLMs generate residue-level embeddings for each sequence in the MSA, the sequence embedding can be directly obtained by aggregating all the residue embeddings in the sequence. Thus we can calculate the cosine similarity matrix between each sequence to measure their pairwise similarities.

To be more specific, we specify two MSA pairing strategies, i.e., Intra-ranking (IntraCos) and Inter-pairing, based on the cosine similarity measurement between sequence embeddings as follows:

#### Intra-ranking (IntraCos)

Firstly, for all sequences from a given MSA *M ∈ 𝒜*^*N×C*^, we obtain a collection of residue-level embedding *{E*_*ln*_ ∈ ℝ^*C×d*^ : *l ∈* [*L*], *n ∈* [*N*]*}*, where *d* is the embedding dimension. For sequence *n ∈* [*N*], we can obtain its sequence-level embeddings *E*_*n*_ = AGG_*l∈*[*L*],*c∈*[*C*]_(*E*_*lnc*_) by aggregating over all layers *L* and all residues *C*, where *E*_*n*_ ∈ ℝ^*d*^. Then we compute cosine similarities between the query sequence embedding, *E*_1_, and other sequence embeddings, *E*_*n*_, where *n* ≠ 1, in the MSA to obtain the pairwise similarity score matrix (IntraCosScore) *S*_1_ ∈ ℝ^1*×N*^. After that, we build interologs like ESMPair does.

##### Inter-ranking

Instead of ranking sequences in each MSA and matching sequences of the same rank, here we directly compute the similarity score matrix between sequences from different MSAs. Formally, given two MSAs 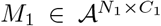 and 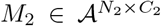, we obtain two individual collections of sequence embeddings 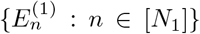 and 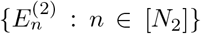. The inter-chain cosine similarity matrix is denoted by 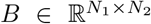, where *B*_*ij*_ = cos(*E*_1_[*i*], *E*_2_[*j*]). Without loss of generality, we assume *N*_*i*_ *≤N*_*j*_, we propose two algorithms to build interologs as follows:

1. **Global Maximization Optimization (InterGlobalCos)**. We formalize the pairing problem as a maximum-weighted bipartite matching problem. The weighted bipartite *G* = (*V, E*) is constructed as follows: sequences from individual MSAs of two chains form the set of vertices in *G*, i.e., 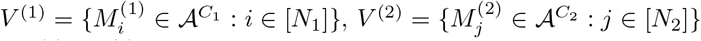, and *V* = *V*^(1)^ ⋃ *V*^(2)^. There are no edges among sequences from the same chain MSA, thus *V*^(1)^ and *V*^(2)^ are two independent sets. There is an edge *e*_*ij*_ between 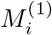 and 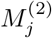 if these two sequences are from the same species; the weight associated with *e*_*ij*_ is *B*_*ij*_. An optimal MSA matching pattern can be obtained by Kuhn-Munkres (KM) algorithm[53] in the polynomial time.
2. **Local Maximization Optimization (InterLocalCos)**. KM algorithm finds a global optimal solution. However, as suggested by [54], in each species, the sequence that is most similar to the query sequence may be more informative, while other sequences that are less similar may add noises. Thus we propose a greedy algorithm that focuses on pairs that have high similarity scores with the query sequence. We iteratively select a pair of sequences (*i, j*) that have the largest score in *B* among sequences that have not been selected before until reaching a pre-defined maximal number of pairs.

#### Complex structure prediction of heteromers with more than two different chains

The proposed methods, such as ESMPair and IntraCos, can be easily extended to build MSA of interologs for heteromers with more than two different chains. In practice, we can rank the MSAs in each query sequence by the similarity matrix obtained by the corresponding metric, then we match them of the same rank in each species to build effective interologs.

### 3.2 Settings

#### Evaluation metric

We evaluate the accuracy of predicted complex structures using DockQ [55], a widely-used metric in the computational structural biology community. Specifically, for each protein complex target, we calculate the highest DockQ score among its top-*N* predicted models selected by their predicted confidences from Alphafold-Multimer. We refer to this metric as the best DockQ among top-*N* predictions.

#### Datasets

In order to investigate how improving pairing MSAs can improve the performance of AlphaFold-Multimer, we construct a test set satisfying the following criteria:

1. There are at least 100 sequences that can be paired given the species constraints.
2. The two constituent chains of a heterodimeric target share *<* 90% sequence identity.

We select heterodimers consisting of chains with 20 ~ 1024 residues (due to the constraint of ESM-MSA-1b and also ignore peptide-protein complex), and the overall number of residues in a dimer is less than 1600 (due to GPU memory constraint) from Protein Data Bank (PDB), as accessed on March 3, 2022. We use the default AlphaFold-Multimer MSA search setting to search the UniProt database [32] with JackHMMER [33], which is used for MSA pairing. We also search the Uniclust30 database [56] with HHblits [57], which is used for monomers, i.e., block diagonal pairing. We further select those heterodimers with at least 100 sequences that can be paired by AlphaFold-Multimer’s default pairing strategy. We define two dimers as at most *x*% similar, if the maximum sequence identity between their constituent monomers is no more than *x*%. Overall, we select 801 heterodimeric targets from PDB that are at most 40% similar to any other targets in the dataset, and satisfy the aforementioned two criteria. Then we use AlphaFold-Multimer (using the default MSA matching algorithm) to predict their complex structures. Based on their predicted confidence scores (pConf) or DockQ scores, 92 targets with their pConf less than 0.7 are denoted as the pConf70 test set. We select 0.7 as the low confidence cutoff based on our fitted logistic regression models over 7,000 DockQ and pConf pairs, because the conditional probability of the model having medium or better quality given pConf equals 0.7 is slightly greater than 0.5 (around 0.6), while the probability is less than 0.5 if pConf equals 0.6. For more comparisons, we also select 0.8 as the cutoff, which results in the pConf80 test set of 168 targets, and 155 targets with their predicted DockQ scores less than 0.49 are denoted as the DockQ49 test set.

#### Baselines

Several heuristic MSA pairing strategies have been developed for protein complex contact and 3D structure prediction [17, 23].

##### Phylogeny-based method

The strategy is first proposed in ComplexContact [28] for complex contact prediction and is widely adopted by the community. AlphaFold-Multimer employed a similar strategy. This strategy first groups sequences in an MSA by their species and then ranks sequences of the same species by their similarity to the query sequence. When there is more than one sequence in a species group, it joins two sequences of the same rank within the same species group to form an interolog. AlphaFold-Multimer uses this strategy and shows state-of-the-art accuracy in complex structure prediction [25]. Practically, we run the implementation code of Alphafold-Multimer following the default setting of official repertory^1^. Notably, we only evaluate the unrelaxed model without the template information for the time efficiency[16].

##### Genetic distances

In bacteria, interacting genes sometimes are co-located in operons and co-transcribed to form protein complexes [58]. Consequently, we can detect interologs by the genetic distance of two genes. This strategy pairs sequences of the same species based on the distances of their positions in the contigs, which are retrieved from ENA. In our implementations, given a sequence from the first chain, we pair it with the sequence from the second chain that is closest to it in terms of genetic distance. If there are more than one closest sequence, we select the one that has the lowest e-value to the query sequence of the second chain; the e-value is calculated by the MSA search algorithm used to construct the chain MSA.

##### Block diagonalization

This strategy pads each chain sequence with gaps to the full length of the complex [23]. Therefore, each sequence in the constructed joint MSA, except for the query sequence, will include non-gap tokens in exactly one chain and gap tokens in other chains. By sorting sequences in the joint MSA, we can make non-gap tokens to appear only in the diagonal blocks, thus this strategy is termed as block diagonalization. In our implementations, given a sequence from the first (second) chain, we append (prepend) non-gap tokens to it until the number of non-gap tokens equals the length of the second (first) chain.

#### Running environment

We conduct the experiments on an Enterprise Linux Server with 56 Intel(R) Xeon(R) Gold 5120 CPU @ 2.20GHz, and a single NVIDIA Tesla V100 SXM2 with 32GB memory size.

## 4 Conclusion & Discussion

This paper explores a series of simple yet effective MSA pairing algorithms based on pre-trained protein language models (PLMs) for constructing effective interologs. To our best knowledge, this is the first time that PLMs are used to construct joint MSAs. Experimental results have confirmed the proposed ESM-Pair significantly outperforms the state-of-the-art phylogeny-based protocol adopted by AlphaFold-Multimer. What’s more, ESMPair performs particularly better on targets from eukaryotes which are hard to be predicted accurately by AF-Multimer. We further confirm that, instead of using the conventional single strategy to build interologs, the ensemble MSA pairing strategy can largely improve the structure prediction accuracy. Generally, ESMPair has a profound impact on biological applications depending on the high-quilty MSA. In the future, we will continue to explore more potential ways to leverage the advantages of PLM in building and choosing MSA. We also looking forward to applying our proposed methods to improve current MSA-based applications.

### Limitations

In this paper, we merely consider how to build effective interologs for heterodimers, which broadly benefits biological applications depending on the high-quality MSA, such as the complex contact prediction [59, 60], complex structure prediction discussed in this paper, etc. However, there also have a large proportion of homodimers in biological assemblies. As it is trivial to build interologs for them, how to select high-quality MSA for homodimers is a more challenging yet important question. Previous work [39, 54] has an empirical insight that instead of using the full MSA searched from the protein sequence database, we can select a few high-quality MSA following some promisings, such as using the MSA maximizing the sequence diversity [39], or choosing the MSA owning the largest sequence similarity with the primary sequence [54]. To date, few efforts have systematically investigated the MSA-selection problem. We leave this for future work.

As we propose a series of MSA paring methods built on the output of PLMs, the representation ability of the PLMs directly affects the performance of our proposed methods. In this paper, we choose the state-of-the-art protein language model so far, i.e., ESM-MSA-1b [39], to support our algorithms. However, it is always worth exploiting the potential correlations between different PLM configurations and the performance of our proposed PLM-enhanced methods to identify effective interologs.

Although ESMPair has advantages over the default strategy adopted by AF-Multimer in identifying MSA of interologs, their success rate is similar. After a deep analysis, we observe ESMPair outperforms AF-Multimer most in acceptable cases (DockQ ≥ 0.23), however it is notoriously difficult for ESMPair to improve DockQ score of unacceptable cases to be acceptable (Only 3% targets). As we follow the pipeline of the complex structure prediction via AF-Multimer (Fig. 1), thus the limited ability of AF-Mulitmer becomes the bottleneck of the performance of ESMPair. Nevertheless, the above extensive experimental results have proved ESMPair consistently outperforms AF-Multimer despite AF-Multimer having an inductive training bias towards its default MSA pairing strategy. From the training process of AF-Multimer, we know that the performance of structure prediction highly depends on the quality of the input MSA. In light of this, we assume that if AF-Muiltimer can fine-tune, or totally train from scratch based on ESMPair’s MSA pairing method, the accuracy of structure predictions may be further improved. Moreover, compared with the conventional MSA pairing method that only uses a single strategy to identify interologs, the ensemble strategy has shown superior performance both in DockQ score and Success Rate without fine-tuning AF-Multimer. We assure that the ensemble strategy proposes a new perspective on how to comprehensively exploit the co-evolutionary patterns among MSA, thus further having a wide impact on the biological algorithms resorting to the input MSA.

## 5 Author Contributions

B.C proposed the main idea, conducted the main experiments, and wrote initial manuscript. Z.W.X, J.Z.Q, and Z.F.Y collected the experimental data, designed experiments, and wrote the initial manuscript. J.B.X and J.T gave the detailed instructions and refined the manuscript.

## 6 Data Availability

Data that involved in this work can be obtained from Github: https://github.com/allanchen95/ESMPair.

## 7 Code Availability

The code of this study can be obtained from GitHub: https://github.com/allanchen95/ESMPair.

---

**Fig. A1.**
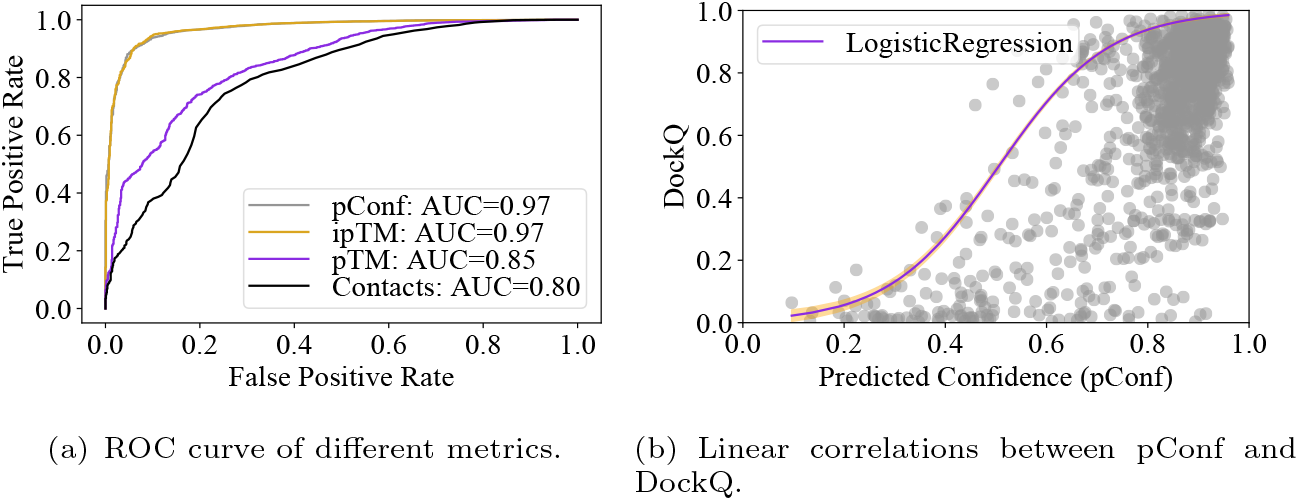
Different metrics assessment. **a**.ROC curve of different metrics of distinguish acceptable cases (DockQ ≥ 0.23) predicted by ESMPair. **b**.The distribution of predicted confidences (pConf, x-axis) and DockQ scores (left y-axis). And the conditional probability of the prediction having DockQ ≥ 0.23 given pConf. The red curve is the visualization of the fitted logistic regression model.

**Fig. A2.**
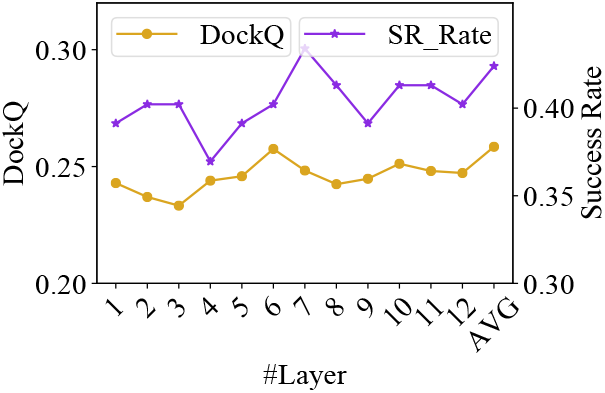
The average of Top-5 Best DockQ scores of ESMPair based on the different layers of ESM-MSA-1b on the pConf70 dataset. AVG means that ESMPair is based on the column-wise attention matrix by averaging the one generated from all the twelve transformer layers.

## Appendix A Supplement Material

### The Number of Effective Interlogs (Meff)

It counts the number of non-redundant interlogs in an MSA, which measures the amount of homologous information. Here we use the toolkit from RaptorX^2^ to estimate the value of Meff. Specifically, we set 70% sequence identity as the cutoff to judge if two interlogs are redundant or not. If the number of interlogs (including itself) similar to interlog *i* is *n*_*i*_, then the weight of interlog *i* is 1*/n*_*i*_. Finally, Meff is calucated by summing the weight of all interlogs.

### Supplement Experiments

We conduct some additionally experiments listed here.

**Table A1.**
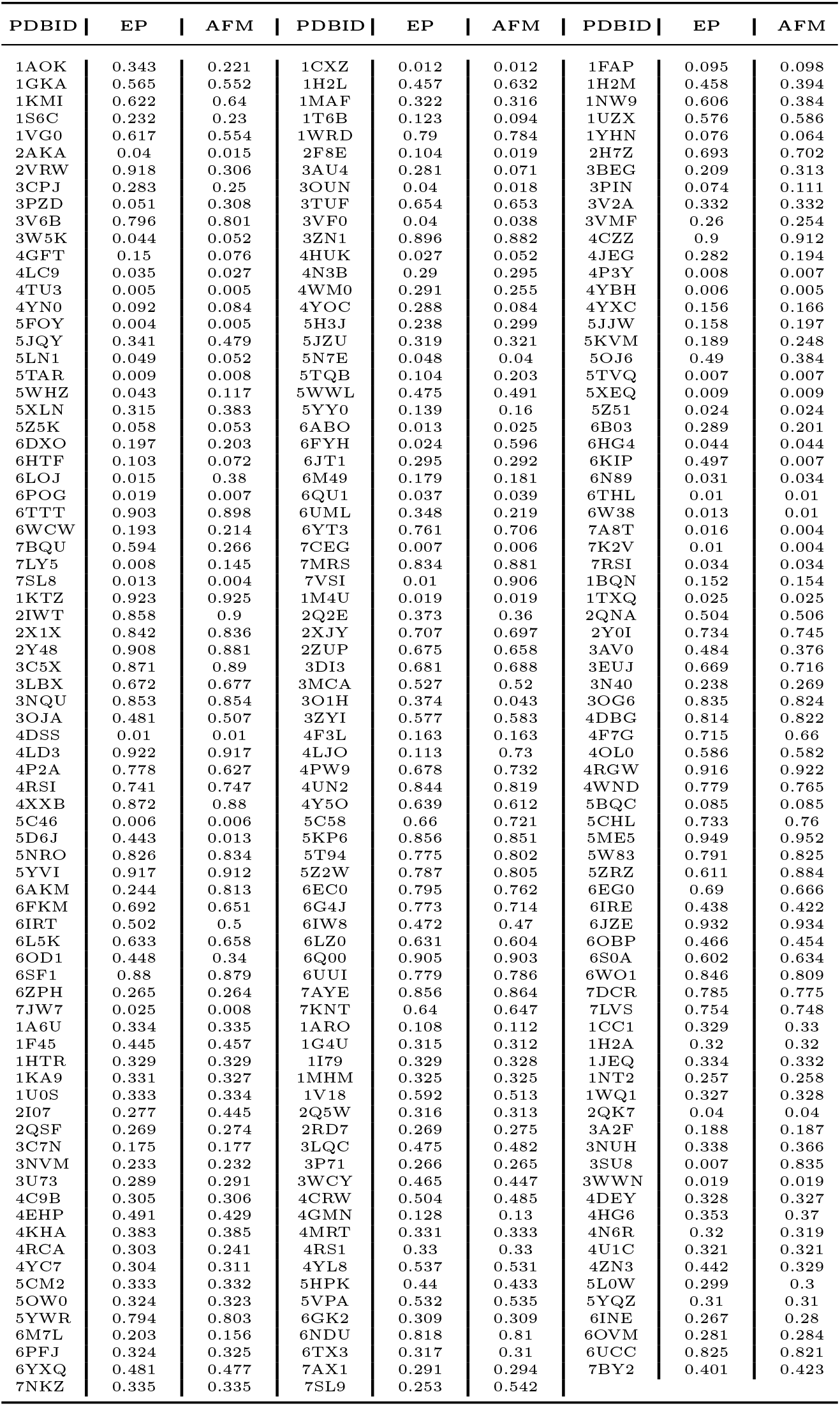
The comparisons of Top-5 Best DockQ scores between ESMPair (EP) and AF-Multimer (AFM) on all test cases.

## Footnotes

1 https://github.com/deepmind/alphafold

2 https://github.com/j3xugit/RaptorX-3DModeling

